# A critical assessment of single-cell transcriptomes sampled following patch-clamp electrophysiology

**DOI:** 10.1101/298133

**Authors:** Shreejoy J. Tripathy, Lilah Toker, Claire Bomkamp, B. Ogan Mancarci, Manuel Belmadani, Paul Pavlidis

## Abstract

Patch-seq, combining patch-clamp electrophysiology with single-cell RNA-sequencing (scRNAseq), enables unprecedented single-cell access to a neuron’s transcriptomic, electrophysiological, and morphological features. Here, we present a systematic review and re-analysis of scRNAseq profiles from 4 recent patch-seq datasets, benchmarking these against analogous profiles from cellular-dissociation based scRNAseq. We found an increased likelihood for off-target cell-type mRNA contamination in patch-seq, likely due to the passage of the patch-pipette through the processes of adjacent cells. We also observed that patch-seq samples varied considerably in the amount of mRNA that could be extracted from each cell, strongly biasing the numbers of detectable genes. We present a straightforward marker gene-based approach for controlling for these artifacts and show that our method improves the correspondence between gene expression and electrophysiological features. Our analysis suggests that these technical confounds likely limit the interpretability of patch-seq based single-cell transcriptomes. However, we provide concrete recommendations for quality control steps that can be performed prior to costly RNA-sequencing to optimize the yield of high quality samples.

## Introduction

Linking gene expression to a neuron’s electrical and morphological features has long been a goal of cellular neuroscience. To this end, one strategy is to use the same patch-clamp electrode for electrophysiological characterization for mRNA sampling, for example, by aspirating the cell’s cytosol into the patch-pipette (Eberwine et al., 1992; Sucher and Deitcher, 1995; Toledo-Rodriguez et al., 2004; Toledo-Rodriguez and Markram, 2014; Kodama et al., 2012; Rossier et al., 2014). The aspirated mRNA transcripts can then be detected and quantified using RT-PCR (Eberwine et al., 1992; Sucher and Deitcher, 1995; Cauli et al., 1997; Toledo-Rodriguez et al., 2004; Kodama et al., 2012; Rossier et al., 2014) or other methods (Subkhankulova et al., 2010), allowing the quantification of multiple genes or transcripts.

Recently, a number of groups have published protocols for patch-seq that extend previous RT-PCR-based methods by quantifying patch-pipette sampled cellular mRNA transcripts using next-generation RNA-sequencing (Cadwell et al., 2015; Fuzik et al., 2016; Földy et al., 2016; Bardy et al., 2016; Cadwell et al., 2017b, 2017a). These protocols make use of recent technical improvements in single-cell RNA-sequencing (scRNAseq) that enable gene expression quantification from very low starting volumes of mRNA (Poulin et al., 2016; Tasic et al., 2017), such as those present in a single-cell or single-nucleus.

Patch-seq mRNA sample collection differs from standard single-cell or single-nucleus RNAseq, in two major ways (Cadwell et al., 2017b, 2017a). First, as opposed to relying on dissociating cells into suspension, the micropipette used for electrical recording is used for mRNA extraction via aspiration. While guiding the patch pipette to (or from) the soma of a cell of interest, the pipette often must travel through the processes of other cells, presenting an opportunity for contamination. Second, the effectiveness of cell content aspiration is difficult to control, so the amount of mRNA extracted may tend to vary from cell to cell.

Here, our goal was to investigate the quality of scRNAseq data profiled using patch-seq. Our strategy was to compare patch-seq derived scRNAseq data with analogous data sampled using cellular-dissociation based methods, from which multiple large and high-quality single-cell transcriptomic datasets are available (Tasic et al., 2016; Zeisel et al., 2015). Our findings suggest that sampling cellular mRNA using a patch-pipette induces technical artifacts that tend not to be present to the same degree in cellular-dissociation based scRNAseq data. Based on our findings, we provide approaches for detecting these technical issues and discuss strategies for generating high-quality patch-seq datasets in the future.

## Methods

### Dataset overview

We made use of 4 previously published patch-seq datasets (Cadwell, Földy, Fuzik, Bardy) (Bardy et al., 2016; Cadwell et al., 2015; Földy et al., 2016; Fuzik et al., 2016), reflecting, to our knowledge, all of the published patch-seq datasets as of January 2018. We compared these to 2 cellular dissociation-based single-cell RNAseq datasets (Tasic, Zeisel) (Tasic et al., 2016; Zeisel et al., 2015). We downloaded single-cell transcriptomic data from each study from accessions provided in Table 1 and Supplementary Table 1 or by contacting the authors directly. We obtained patch-seq-based electrophysiological data for the Cadwell and Fuzik datasets from the authors. For all patch-seq datasets, electrophysiological data were provided as a spreadsheet containing a set of summarized electrophysiological features per cell (e.g., input resistance, resting membrane potential, etc.). Electrophysiological data from the Allen Institute Cell Types database (celltypes.brain-map.org) were obtained and processed as described previously (Tripathy et al., 2017).

### Transcriptome data pre-processing

We reprocessed transcriptomic data for the Cadwell, Földy, and Tasic datasets directly from Gene Expression Omnibus (GEO) or Array Express. Data from GEO was downloaded using fastq-dump version 2.8.2 from the Sequence Read Archive Toolkit. Technical reads such as barcodes and primers were filtered out during extraction. Adapter sequences were clipped from the raw reads. The list of option used is as follows: ‘--gzip --skip-technical --readids --dumpbase --split-files --clip’. Data from ArrayExpress was downloaded and used directly as prepared by the European Bioinformatics Institute.

The reference mouse transcriptome was produced using the ‘rsem-prepare-reference’ script provided by the RSEM RNA-Seq transcript quantifier (Li and Dewey, 2011). The assembly version used was Ensembl GRCm38, packaged by Illumina for the iGenomes collection. Alignment was performed using STAR (Dobin et al., 2013) version 2.4.0h, provided as the aligner to RSEM v1.2.31. Default parameters were used (with the exception of parallel processing and logging related options). Transcript definitions used to detect ERCC spike-ins were obtained from the ERCC92 version fasta and GTF files. Spike-ins were concatenated to the GRCm38 assembly before applying rsem-prepare-reference, and independently to create a standalone ERCC assembly. Both the concatenated and standalone spike-ins assemblies showed highly comparable proportions of spike-in expression. For the Fuzik and Zeisel datasets, we made use of the quantified summarized unique molecule counts (UMIs) made available at GEO. For the Bardy dataset, we used the summarized count matrices directly provided by the authors.

### Mapping of mouse patch-seq cell types onto taxonomies derived from dissociated cells

Using descriptions for cellular identities provided in the original patch-seq publications, we manually mapped each of the cell types represented across the three mouse patch-seq datasets onto transcriptomically-defined cellular clusters reported in the two dissociated cell datasets (shown in Supplementary Table 2). For example, given that the elongated neurogliaform cells and single bouquet cells characterized in Cadwell are both cortical layer 1 cells, we manually mapped these to the layer 1 cells defined in Tasic as Ndnf cells. Similarly, we mapped the hippocampal regular-spiking interneurons characterized in Foldy to the Sncg cluster from Tasic (personal communication with Csaba Földy). To align cell subtype clusters between Tasic and Zeisel, we used mappings provided by MetaNeighbor (Crow et al., 2018) (shown in Supplementary Table 2). The mappings between broad cell types in Tasic with Zeisel are provided in Supplementary Table 3. As with our previous work mapping cells and cell types across datasets (Mancarci et al., 2017; Tripathy et al., 2017), we note that these cross-dataset mappings are approximate and ideally would be guided by the use methods for unambiguously aligning cell types across experiments (e.g., transgenic mouse lines with specific cell types labeled by fluorescent proteins).

### Identification of cell type-specific marker genes

For this study, we defined two classes of marker genes, termed “on” and “off” markers. The first class, “on” markers, are genes that are highly and ubiquitously expressed in the cell type of interest with enriched expression relative to other cell types. The second class, “off” markers, are expected to be expressed at low levels in a given patch-seq cell type. These are genes that are specifically expressed in a single cell type (e.g., microglia) and, if expressed, are an indicator of possible cellular contamination. To identify marker genes, we employed two recent surveys of mouse cortical diversity from Tasic et al. and Zeisel et al. (Tasic et al., 2016; Zeisel et al., 2015).

To identify “on” marker genes, we initially used the Tasic dataset, and selected genes whose average expression in the chosen cell type was >10 times relative all other cell types in the dataset, with an average expression in the cell type of >100 TPM. From this initial gene list, we next filtered these genes to only include those that were expressed >10 TPM/cell in >75% of all cells of that type in Tasic, and >1 UMI/cell in >50% of all cells of that type in Zeisel. Using the Tasic nomenclature, we defined “on” markers for Ndnf, Sncg, Pvalb, and Pyramidal cell types.

To identify “off” marker genes for broad cell types (shown in Supplementary Table 3), as an initial listing we used the set of cell type-specific marker genes for broad cell classes in the mouse cortex, defined in our previous work using the NeuroExpresso database (Tasic et al., 2016). Specifically, we used the set of cortical markers derived from single-cell RNA-seq for astrocytes, endothelial cells, microglia, oligodendrocytes, oligodendrocyte precursor cells, and pyramidal cells. From this list, we first filtered out lowly expressed genes that were expressed <10 TPM/cell in >50% of all cells of that type in Tasic, and <1 UMI/cell in >50% of all cells of that type in Zeisel. Next, we filtered genes too broadly expressed in our patch-seq cell types of interest by assessing the expression of these genes in the Ndnf, Sncg, Pvalb, and Pyramidal cell types, removing genes that were expressed at a level greater than >10 TPM/cell in >33% of all cells of that type in Tasic, and >2 UMI/cell in >33% of all cells of that type in Zeisel.

When defining on and off marker genes for inhibitory cell subtypes (e.g., the Ndnf cell type), we did not compare these cells to other GABAergic cells. For example, when defining “on” markers for Ndnf cells, we did not compare these cells’ expression to Pvalb or Sst cells. We note that this choice limits our ability to identify inhibitory-to-inhibitory cell contamination, for example, an Ndnf cell contaminated by Sst-cell specific markers. To define an initial set of “off” markers for GABAergic inhibitory cells, we first obtained a list of genes based on Tasic where in GABAergic cells had average expression >10 times all other non-GABAergic cells in the dataset and with an average expression of at least 100 TPM.

The final list of filtered mouse cell type specific marker genes used in this study are provided in Supplementary Table 4.

To obtain a list of human cell type specific marker genes for use for the Bardy dataset, we made use of classic cell-type specific markers for astrocytes and microglia, based on human purified cell types shown in Figure 4A of reference (Zhang et al., 2016).

### Summarizing cell type-specific marker expression

When directly comparing expression values from patch-seq data to dissociated cell data, we compared the Cadwell and Földy datasets to Tasic, as these all were quantified using TPM and employed Smart-seq-based methods. Similarly, we compared Fuzik dataset to Zeisel, as these both used C1-STRT and were quantified using unique molecule identifiers (UMIs), normalized as UMI counts per million. We summarized a single-cell sample’s expression of multiple cell type-specific markers using the sum of the log_2_ normalized expression values. Given a patch-seq sample of cell type identity A (e.g., a pyramidal cell) and wanting to quantify its normalized expression of “off” markers for cell type B (e.g., microglial markers), we used the dissociated cell data to estimate the median expression of cell type B’s “markers in cells of type A (e.g., median expression level of microglial markers in pyramidal cells) and the median expression of cell type B’s markers in cells of type B (e.g., median expression level of microglial markers in microglia cells). Specifically, we normalized expression to a value of approximately 0 to 1, as follows:

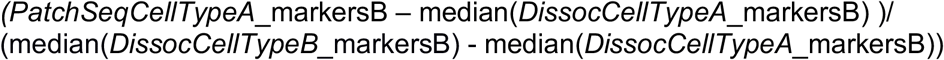

where we set all negative values to 0. Next, to obtain a single contamination index per single cell, we summed all contamination scores for all broad cell types, excluding the patch-seq cell’s assigned broad cell type.

Lastly, to obtain a scalar quality score for transcriptomic data from patch-seq samples (e.g., for analysis of electrophysiological data), we used the Spearman correlation of each patch-seq sample’s expression of “on” and “off” marker genes to the average expression profile of dissociated cells of the same cell type (shown in Supplement Figure 3). For example, for an Ndnf patch-seq sample from Cadwell, we first calculated the average expression profile of Ndnf cells from Tasic across the set of all “on” and “off” marker genes (i.e., Ndnf markers, pyramidal cell markers, astrocyte markers, etc.), and then calculated the correlation between the patch-seq cell’s marker expression to the mean dissociated cell expression profile. Since these correlations could potentially be negative, we set quality scores to a minimum of 0.1. A convenient feature of this quality score is that it yields low correlations for samples with relatively high contamination as well as those where contamination is largely undetected but expression of endogenous “on” markers is also low (Supplement Figure 3).

### Analysis of factors influencing the numbers of genes detected per cell

We analyzed how the following factors influenced the numbers of genes detected per cell: library size, defined as the total numbers of reads sequenced per cell; spike-in ratio, defined as the number of reads mapping to ERCC spike-ins divided by total sequenced reads; the unmapped ratio, defined as the ratio of reads not mapping to the exonic reference divided by all non-ERCC sequenced reads; and cellular contamination indices, as defined in the previous section. For the Cadwell, Tasic, and ERCC-containing subsets of the Földy, and Bardy datasets, we fit a linear model (implemented using the ‘lm’ function in R) for numbers of detected genes per each cell as follows:

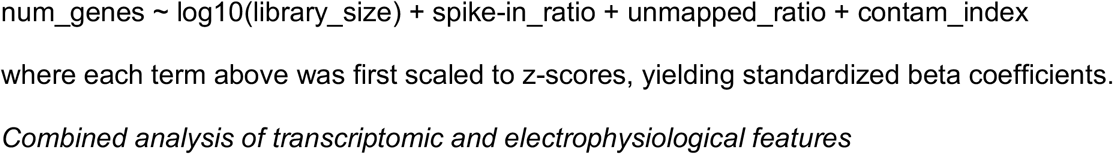

where each term above was first scaled to z-scores, yielding standardized beta coefficients.

### Combined analysis of transcriptomic and electrophysiological features

We analyzed correlations between transcriptomic and electrophysiological features using an approach similar to our previous work (Tripathy et al., 2017). For each patch-seq dataset, we first filtered for genes whose average expression was > 30^th^ percentile relative to all genes in the dataset. We analyzed electrophysiological features overlapping with our previous analysis, specifically, input resistance (Rin), resting membrane potential (Vrest, action potential threshold (APthr), action potential amplitude (APamp), action potential half-width (APhw), membrane time constant (Tau), after-hyperpolarization amplitude (AHPamp), rheobase (Rheo), maximum firing rate (FRmax), and capacitance (Cm). We calculated Pearson correlations between the set of electrophysiology features and gene expression values, both without weighting cells by their overall quality scores (based on correlation of markers to dissociated cell samples), and after weighting cells using their quality scores.

We performed an analogous analysis for comparison of pooled-cell correlations based on the AIBS/Tasic dataset, where we computationally merged different groups of cells characterized using dissociated cell scRNAseq (based on Tasic et al, (Tasic et al., 2016)) with cells characterized using patch-clamp electrophysiology (Teeter et al., 2018) based on the overlap of same mouse transgenic lines and coarse cortical layers (i.e., upper vs lower mouse visual cortex). For example, we merged 14 QC-passing scRNAseq samples from the Sst-IRES-cre mouse line from visual cortex dissections specific to lower layers with 89 patch-clamp samples from the same mouse line from cortical layers 4 through 6b. After merging single-cells into cell types, we averaged expression and electrophysiological values; since cell types tended to be represented by differing numbers of cells, in our gene-electrophysiology correlation analyses we weighted cell types based on the numbers of cells available using the square root of the harmonic mean of the number of cells characterized by electrophysiology and electrophysiology.

### Statistical information

We used the R weights toolbox (v0.85) to calculate weighted Pearson correlations and raw p-values. We used the Benjamini-Hochberg False Discovery Rate (FDR) to account for analysis of multiple correlations.

### Computer code and data availability

All computational code and associated data has been made accessible at https://github.com/PavlidisLab/patchSeqQC and code for the RNAseq pipeline is accessible at https://github.com/PavlidisLab/rnaseq-pipeline.

## Results

To quantitatively assess the influence of patch-seq specific technical confounds, we performed a re-analysis of four recently published patch-seq datasets. We focused our analyses on three datasets obtained from mouse acute brain slices (Cadwell et al., 2015; Földy et al., 2016; Fuzik et al., 2016) and contrast these against one dataset obtained from human stem-cell derived neurons and astrocytes in culture (Bardy et al., 2016) (Table 1).

**Table 1:** Description of patch-seq datasets re-analyzed in this study. *Expression data obtained by contacting the authors directly.

| Dataset | Description | Preparation | RNA amplification | Number of cells | Accession |
| --- | --- | --- | --- | --- | --- |
| Cadwell (Cadwell et al., 2015) | Cortical layer 1 interneurons | Acute mouse slices | Smart-seq2 | 58 | E-MTAB-4092 |
| Fuzik (Fuzik et al., 2016) | Cortical layer 1/2 interneurons and pyramidal cells | Acute mouse slices | STRT-C1 (with unique molecule identifiers) | 80 | GSE70844 |
| Földy (Földy et al., 2016) | Hippocampal CA1 and Subiculum pyramidal cells and regular- and fast-spiking interneurons | Acute mouse slices | SMARTer | 93 | GSE75386 |
| Bardy (Bardy et al., 2016) | Stem-cell derived neurons and astrocytes | Differentiated human cells in culture | SMARTer | 56 | NA* |

### Expression of off-target cell type marker genes in patch-seq samples

We first assessed if patch-seq based single-cell transcriptomes might have been contaminated by mRNA from other cells adjacent to the patched cell (Figure 1A, B), termed off-target cell-type contamination (Okaty et al., 2011). For example, is there paradoxical expression of genes specific to microglia in the scRNAseq profile of a recorded pyramidal cell? To address this question, we made use of the fact that the broad identities of the recorded cells can be ascertained from morphological and electrophysiological features without relying on the transcriptomic data (see Methods). Furthermore, we used multiple mouse forebrain scRNAseq datasets collected from dissociated cells to define lists of marker genes specific to various cortical and hippocampal cell types (Supplementary Table 4) (Mancarci et al., 2017; Tasic et al., 2016; Zeisel et al., 2015).

We detected that some of the single cell samples from the three mouse datasets collected from acute brain slices expressed markers for multiple distinct cell types (Figure 1, Supplement Figure 1). For example, some of the cortical layer 1 elongated neurogliaform cells (eNGCs) characterized in the Cadwell dataset appeared to also express multiple marker genes specific to pyramidal cells (Figure 1C), such as *Slc17a7*, the vesicular glutamatergic transporter VGLUT1. Similarly, many of the cells identified as hippocampal regular spiking GABAergic interneurons in the Földy dataset also expressed microglial and pyramidal cell markers (Figure 1H).

We sought to quantify the extent of off-target cell type contamination in the mouse patch-seq samples. We directly compared the patch-seq-based expression profiles to cellular dissociation-based transcriptomes from two recent surveys of mouse cortical diversity from Tasic et al. and Zeisel et al. (Tasic et al., 2016; Zeisel et al., 2015). After matching cell type identities across studies (shown in Supplementary Table 2), we found that compared to dissociated cells, patch-seq-based samples expressed markers for multiple cell types at considerably higher levels (Figure 1C, H, J; Supplement Figure 2A, B). We defined a simple contamination index, providing a scalar value for greater than expected off-target cell type marker expression across multiple classes of broad cell types, by comparison to analogous cells from the dissociated-cell reference (see Methods). Importantly, patch-seq-based samples with larger contamination indices also expressed markers of their own cell type at lower levels (Supplement Figure 3). We note that we saw less off-target cell type marker expression in the Fuzik dataset relative to the Cadwell and Földy datasets (Supplement Figure 2), suggesting either less contamination in these cells or that the lower gene detection rate in this dataset (Figure 3B) obscures our ability to use expression profiles to identify cellular contamination.

**figure 1:**
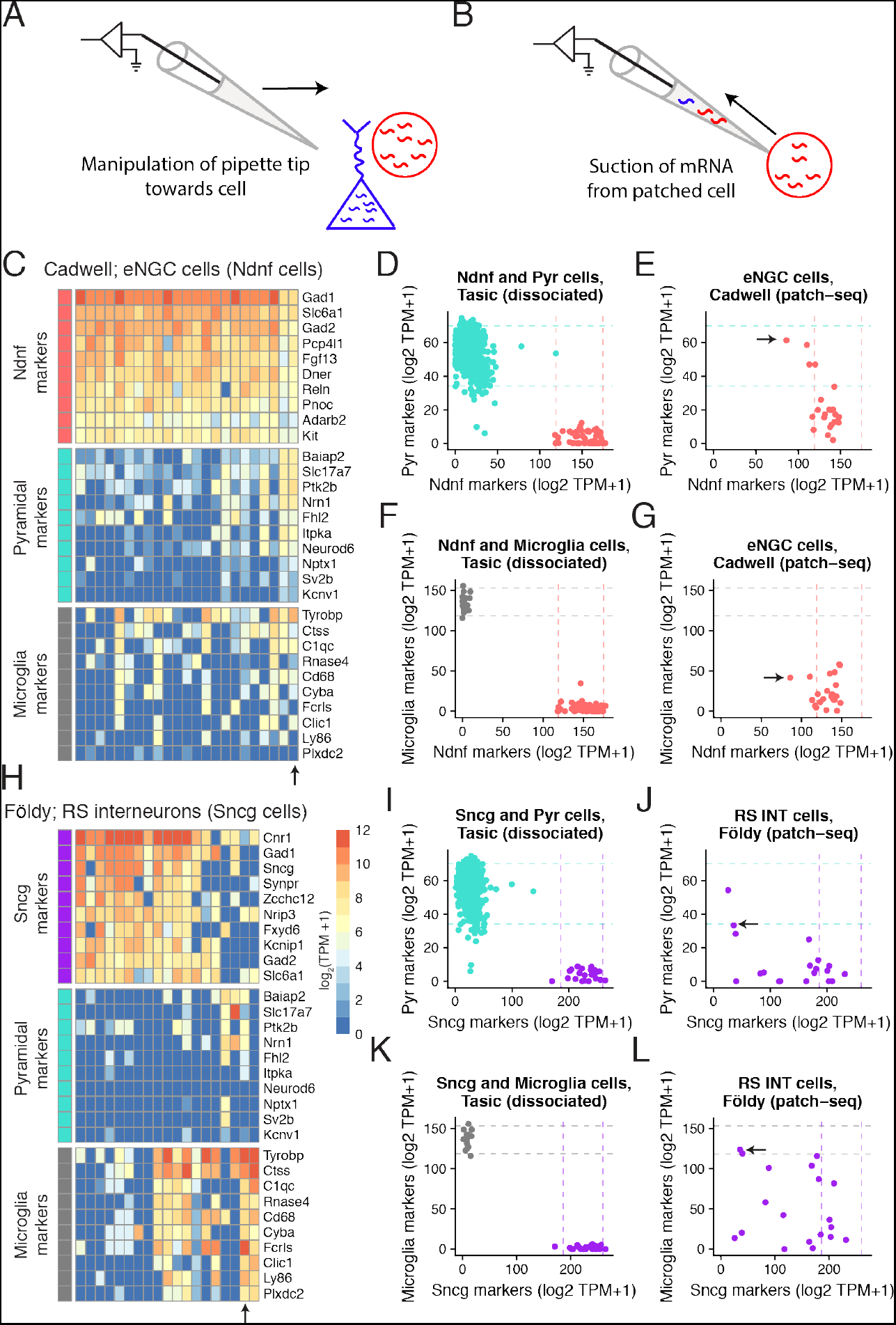
Expression of cell type-specific marker genes in mouse single-cell samples collected using patch-seq. A, B) Schematic illustrating manipulation of patch-pipette towards cell of interest (A) and aspiration of cellular mRNA into the patch-pipette (B). C) Gene expression profiles for GABAergic elongated neurogliaform cells (eNGCs, similar to layer 1 Ndnf cellular subtype) for various cell type-specific markers. Each column reflects a single-cell sample. D) Summed expression of cell type-specific marker genes for Pyramidal cell (y-axis) and Layer 1 Ndnf cell (x-axis) markers. Dots reflect Pyramidal (turquoise) and Ndnf (red) single cells collected in Tasic dataset, based on dissociated scRNAseq. Dashed lines reflect 95% intervals of marker expression for each cell type. E) Same as D, but showing summed marker expression for eNGC cells shown in A based on patch-seq data. Arrow shows single-cell marked in C. F,G) Same as D and E, but for microglial cell markers. H-L) Same as C-G, but for hippocampal GABAergic regular spiking interneurons (RS INT cells, similar to Sncg cells from in Tasic) characterized in Földy dataset.

We next assessed the degree of off-target cell type contamination in the Bardy patch-seq dataset of human stem-cell derived neurons and astrocytes obtained from cultured cells (Bardy et al., 2016). Since the cells in this dataset were cultured relatively sparsely, allowing the processes of each cultured cell to be easily visualized (personal communication with Cedric Bardy), we wondered if this dataset would show less off-target cell type marker expression compared to the three mouse acute brain slice datasets. Indeed, when assessing astrocyte marker expression in the population of electrophysiologically-mature neurons (with markers based on purified human cells (Zhang et al., 2016)), we found these neurons showed some, but overall very little, expression of astrocyte markers relative to the mature astrocytes also profiled in this dataset (Figure 2A, B). In addition, both neurons and astrocytes showed almost no expression of microglia markers (Figure 2A), perhaps unsurprisingly, since microglial cells are not present in these cultures (Bardy et al., 2016). This example provides suggestive evidence that the density of processes of adjacent cells might contribute to off-target mRNA contamination.

**figure 2.**
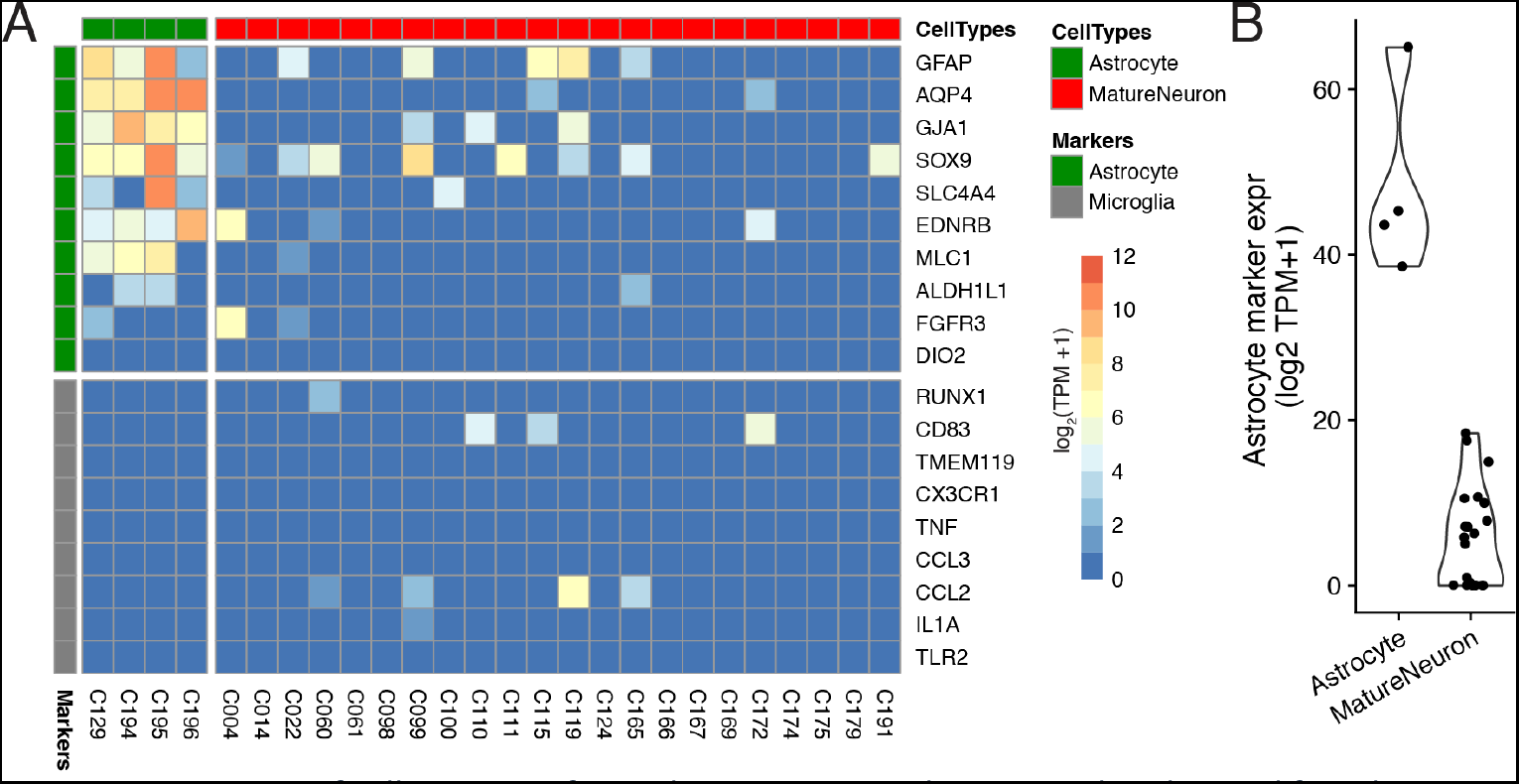
Expression of cell type-specific marker genes in patch-seq samples obtained from human astrocytes and neurons differentiated in culture from the Bardy dataset. A) Gene expression profiles for differentiated astrocytes (green) and electrophysiologically-mature neurons (red) for astrocyte and microglial-specific (grey) marker genes. Each column reflects a single-cell sample. Two astrocyte cells were removed because they expressed fewer than 3 astrocyte markers. B) Summed astrocyte marker expression for astrocyte and mature neuron single-cells, for the same cells shown in part A.

### Technical factors strongly influence the numbers of genes detected per cell

Next, we wondered if there are identifiable technical factors that can help explain the large ranges in the numbers of genes detected per cell in each dataset, from 6000-13000 genes/cell in Cadwell to 800-7000 genes/cell in Fuzik (Figure 3B). Because patch-seq mRNA collection requires the experimenter to manually aspirate cellular mRNA into the patch-pipette, we reasoned that mRNA harvesting would be difficult to consistently control from cell to cell, leading there to be different amounts of extracted mRNA per cell. To estimate how much cellular mRNA was extracted per cell, we made use of ERCC spike-ins (Tasic et al., 2017), which are synthetic control mRNAs that are added to single-cell samples prior to library preparation and sequencing (Figure 3A). Specifically, since the same amount of ERCC spike-in mRNAs are added to each sample, we can use the ratio of spike-in reads to the total count of sequenced reads to estimate the relative amount of extracted mRNA per cell (Lun et al., 2017; Vallejos et al., 2017). Here, every cell in the Cadwell and Tasic datasets and a subset of cells in the Földy and Bardy datasets contained ERCC spike-ins.

We used a multivariate regression approach to ask how various technical factors contribute to the numbers of genes detected per cell in the Cadwell, Földy, and Bardy patch-seq datasets and the Ndnf cell subset of the Tasic dissociated-cell dataset (Figure 3C; the Fuzik dataset did not include spike-ins). Library size (the number of sequenced reads per cell) was positively correlated with detected gene counts in the Tasic and Cadwell datasets (Figure 3C, D). Similarly, cells with a larger ratio of spike-in reads to total sequenced reads (i.e., with lower initial amounts of cellular mRNA; Figure 3A), had lower numbers of detected genes across all of the datasets (Figure 3D), pointing to the importance of mRNA extraction efficiency. In addition, we saw considerably greater ranges in the spike-in ratio in the patch-seq datasets relative to the Tasic dataset (Cadwell: 3-17%, Bardy: 3-37x0025;, Tasic: .4-4%).

Next, we reasoned that though many mRNA transcripts might be extracted from a cell, not all of these would be sufficiently high quality to map to the reference (e.g., they might reflect degraded mRNAs (Cadwell et al., 2017b, 2017a), other contaminants, etc.). To account for this possibility, we calculated the ratio of unmapped to mapped reads, after excluding reads mapping to spike-ins. Cells with very large ratios of unmapped to mapped reads had fewer genes detected (Figure 3C). This technical factor was especially important in the Földy and Bardy datasets, with some cells in the Földy dataset having fewer than 10% of reads mapped to the transcriptome (Figure 3G). Lastly, we further wondered if cells showing greater amounts of off-target cell type contamination would also have a greater number of detected genes. We found that cells with greater contamination indices from the Cadwell and Földy datasets (i.e., the acute slice-based patch-seq datasets) had more genes detected, consistent with previous reports (Ilicic et al., 2016; Vallejos et al., 2017). In total, these simple technical factors explain between 50-85% of the cell-to-cell variance in the detected gene counts per patch-seq datasets (Figure 3I).

**figure 3.**
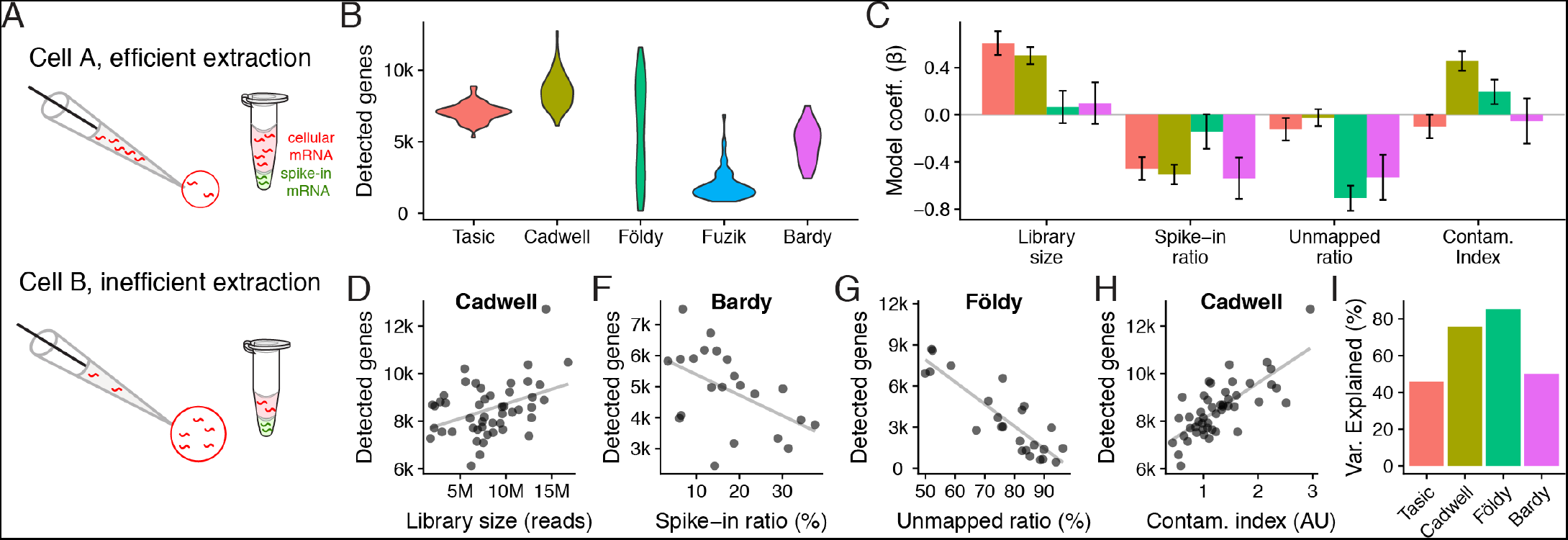
Patch-seq experimental confounds affect the numbers of genes detected per cell. A) Schematic illustrating how spike-in mRNAs can be used to estimate how much mRNA was extracted per cell. B) Violin plots showing numbers of protein-coding genes detected per cell across patch-seq datasets or the Ndnf subset of the Tasic dissociated-cell dataset. C) Technical factors associated with numbers of genes detected per cell across datasets (dataset color shown in B). Bars show standardized beta model coefficients with y-axis in units of standard deviations, allowing comparison of effects across factors and across datasets. Error bars indicate coefficient standard deviations. Positive (negative) coefficients indicate factor is correlated with increased (decreased) gene counts. Regression models calculated using only cells containing mRNA spike-ins. D-H) Examples of univariate relationships between technical factors and detected gene count per cell (dots) across patch-seq datasets. Grey line shows best fit line. D) Library size (count of sequenced reads per cell). F) Spike-ins as a fraction of all sequenced reads per cell. Samples with lower cellular mRNA content (indicated by higher spike-in ratios) have lower gene counts. G) Unmapped ratio, calculated as the ratio of exonic reads to all other reads (excluding spike-ins). H) Cellular contamination index, quantified by summing normalized contamination values across tested cell types (arbitrary units). F) Overall percent variance explained by each dataset-specific statistical model shown in E.

### Accounting for technical factors improves the correspondence with electrophysiological features

Lastly, we performed an integrated analysis of gene expression and electrophysiological features for the 3 mouse-based patch-seq datasets, reasoning that more lower quality patch-seq samples would be less informative of relationships between cellular electrophysiology and gene expression (Tripathy et al., 2017). We first calculated a quality score for each patch-seq sample, based on the similarity of its marker expression to dissociated cells of its same type (see Methods; Supplement Figure 3). After statistically down-weighting lower quality cells, we observed a modest improvement in the correspondence between gene expression and electrophysiology, as evidenced by an increase in the number of genes significantly correlated with electrophysiological features (FDR < 0.1, Figure 4A, B). In addition, after correction, we found more genes overlapping with those identified in our previous gene-electrophysiology correlation analysis based on pooled cell types (Tripathy et al., 2017) (Figure 4C). While the biological implications of these correlations require further investigation, this analysis suggests that controlling for these technical factors can help improve the interpretability of patch-seq data.

**figure 4.**
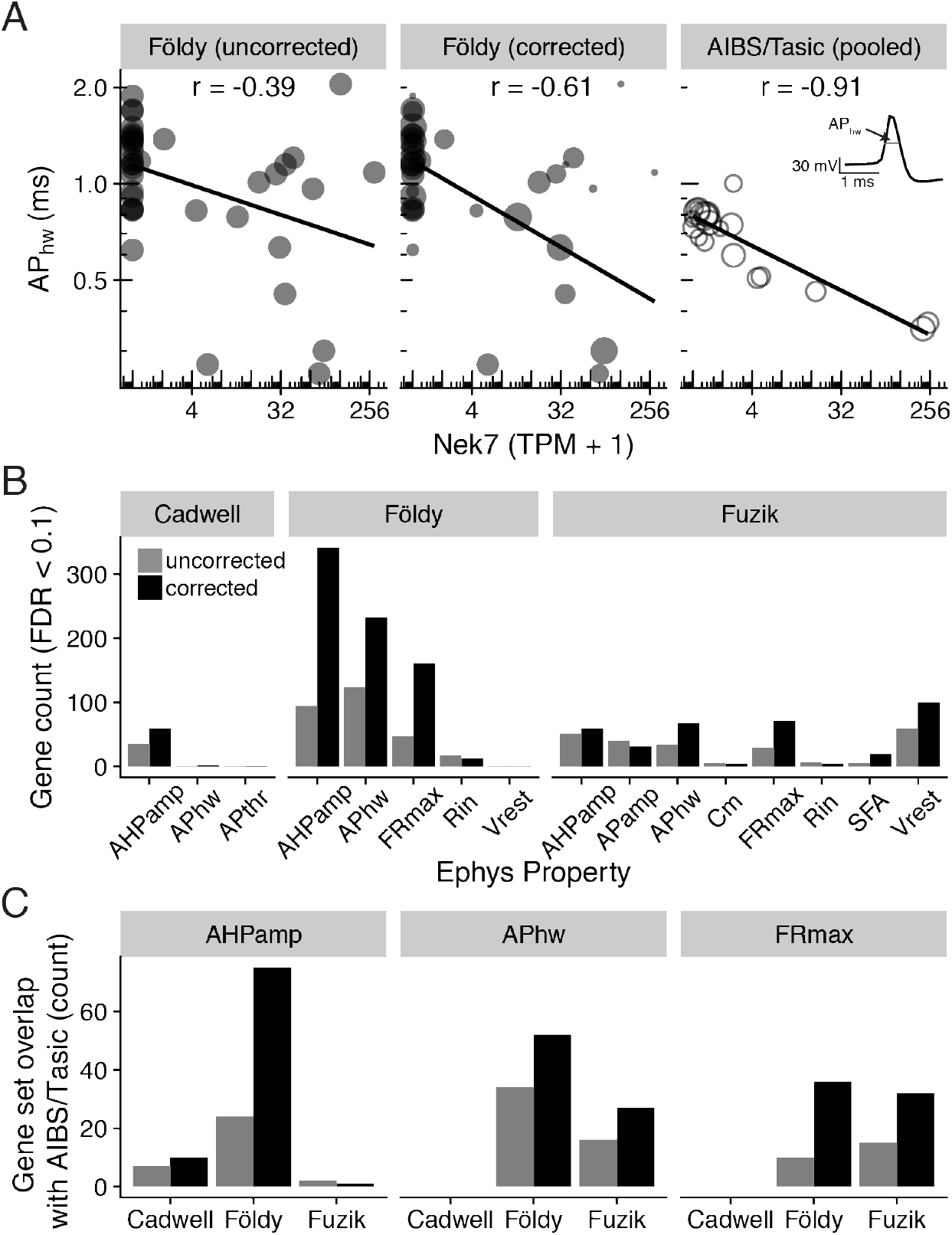
Adjusting for patch-seq experimental confounds improves the correspondence with electrophysiological measures. A) Comparison of gene expression (Nek7; x-axis) with electrophysiological features (action potential half-width; AP_hw_; y-axis). Left panel shows single-cell samples (circles) from the Földy dataset. Middle panel shows same data as left, but size of circles proportional to each sample’s quality score, defined as the similarity of marker expression to dissociated cell-based reference data. Right panel shows cell type-level analysis based on pooled cell type data from Allen Institute cell types database (AIBS/Tasic), where scRNAseq and electrophysiology were performed on different cells from same type (Tripathy et al., 2017). Each open circle reflects one cell type and circle size is proportional to the number of cells representing each cell type. Inset illustrates calculation of action potential half-width (schematic). B) Count of genes significantly correlated (FDR < 0.1) with various electrophysiological properties before (grey) and after (black) correcting for contamination. C) Comparison of genes significantly correlated (BH FDR < 0.1) with electrophysiological features based on patch-seq data with analogous correlations based on AIBS/Tasic dataset, pooled to the level of cell types based on cre-lines. Bars indicate count of overlapping genes between patch-seq and AIBS/Tasic pooled-cell data without correcting for contamination and with correction. No maximum firing rate (FRmax) electrophysiological features were originally calculated for cells in the Cadwell dataset.

## Discussion

The patch-seq technique reflects a considerable leap in our ability to interrogate a neuron across multiple features of its activity. However, across our analyses of multiple patch-seq datasets, we noticed several technical issues that appeared to be shared across experiments. First, in the three mouse datasets collected from acute brain slices (Cadwell et al., 2015; Földy et al., 2016; Fuzik et al., 2016), we observed that many single cell samples appeared to strongly express marker genes from off-target cell types. We interpret this as mRNA contamination from cells adjacent to the recorded cell, but note that there are alternative explanations. Second, we observed that mRNA extraction efficiency differs between sampled cells, leading to varying numbers of genes detected even among cells of the same broad type. These technical artifacts can be mitigated in part through post hoc analyses, such as our attempt to weight single-cells by the similarity of their marker gene expression to analogous dissociated cells of the same broad cell type.

To detect off-target cell type contamination, our main approach was to compare patch-seq based single-cell transcriptomes to dissociated-cell based reference scRNAseq data from similar cell types. We used these reference data to identify cell type-specific marker genes as well as to determine approximately how much off-target marker expression would be expected in each cell type. We note that there are obvious methodological differences between dissociated-cell scRNAseq and patch-seq (Cadwell et al., 2017b, 2017a), such as the strain induced by dissociating cells (Wu et al., 2017) or that patch-seq might be more likely to sample transcripts from distal cellular processes. Thus we cannot conclusively rule out that some of the off-target cell type marker expression might reflect a true biological signal, as opposed to mRNA contamination from adjacent cells. However, we note that the use of marker genes to identify suspected off-target contamination is a routine quality control step in cell type-specific gene expression analyses (Mancarci et al., 2017; Okaty et al., 2011), including recent methods for identifying suspected ȁCdoublets” or multi-cell contamination in droplet-based scRNAseq (Zeisel et al., 2018).

We speculate that the sources of off-target contamination are the processes of cells adjacent to the patch-pipette. For example, while there are relatively few cell bodies in layer 1 of the neocortex, there are processes of other cell types like pyramidal cells, and it is well established that these processes contain mRNA transcripts (Glock et al., 2017). In addition, we noticed that we routinely observed expression of microglial markers in the mouse patch-seq samples. This is interesting because the presence of even 1 mM ATP in the patch-pipette is sufficient to induce rapid chemotaxis of microglial processes towards the pipette (Madry et al., 2018). Patch-clamp intracellular solutions usually use 2 or 4 mM ATP (Tebaykin et al., 2017), including those of the patch-seq datasets here (Bardy et al., 2016; Cadwell et al., 2015; Földy et al., 2016; Fuzik et al., 2016). At present, it is unclear whether this suspected off-target contamination might occur while the pipette is actively manipulated under positive pressure towards the recorded cell.

Alternatively, such contamination might take place following mRNA extraction during the retraction of the pipette from the neuropil and recording chamber. Assuming that neuropil is the major source of off-target contamination, this suggests that there may be advantages to performing patch-seq on sparsely cultured or acutely dissociated cells (Bardy et al., 2016; Kodama et al., 2012; Schulz et al., 2006).

Our analyses identified several technical factors that influence the numbers of genes detected per cell. First, to obtain a sufficient number of detected genes, it is essential to extract a large amount of mRNA from the targeted cell. However, this itself is not sufficient, as other factors, such as mRNA degradation can lead the extracted transcripts being too low quality to map to the genomic reference (Cadwell et al., 2017b, 2017a). Second, given sufficient extraction of non-degraded transcripts, because of the extremely high sensitivity of modern ultra-low mRNA capture kits (Poulin et al., 2016; Tasic et al., 2017), any off-target cell-type contamination will inflate the numbers of genes detected per cell. This suggests that the detected gene count, often used as a proxy for the quality of scRNAseq data, should not be the only quality control metric for single-cell transcriptomes sampled using patch-seq.

The effect of these technical confounds on downstream analyses of patch-seq data is likely context specific. For example, the presence of a small degree of off-target contamination is likely to be of little consequence if the patch-seq data is used as a “Rosetta stone”, to help connect cellular classifications based on different methodologies, such as transcriptomically-defined cell clusters with electrophysiological clusters (Fuzik et al., 2016; Tasic, 2018). However, accurately quantifying single-cell transcriptomes is likely to be much more important when using these data to investigate how transcriptomic heterogeneity gives rise to subtle cell to cell variability in physiological features (Cadwell et al., 2015; Schulz et al., 2006; Tripathy et al., 2017).

Our analyses point to quality control steps that can improve the yield of high-quality patch-seq samples. An advantage of patch-seq over traditional dissociated-cell based scRNA-seq is that a cell’s electrophysiological and morphological features are often sufficient to determine its broad cell type (Cadwell et al., 2015; Földy et al., 2016; Fuzik et al., 2016). We argue that knowing a cell’s broad type can help quality control its sampled transcriptome: the cell should express marker genes of its own type, including highly expressed markers as well as more lowly expressed markers, such as some transcription factors and long non-coding RNAs (Mancarci et al., 2017). In addition, the cell should not express marker genes specific to other cell types. This quality control step can be performed following RNAseq, as we pursue here. However, this quality control could also be performed after library preparation and amplification but prior to costly sequencing, for example, using qPCR to detect the expression of a small number of expected and unexpected marker genes (Bardy et al., 2016).

To summarize, though patch-seq provides a powerful method for multi-modal neuronal characterization (Bardy et al., 2016; Cadwell et al., 2017b; Földy et al., 2016; Fuzik et al., 2016), it is susceptible to a number of methodology-specific technical artifacts, such as an increased likelihood of mRNA contamination from adjacent cells. These artifacts strongly bias traditional scRNAseq quality metrics such as the numbers of genes detected per cell. By leveraging high-quality reference atlases of single-cell transcriptomic diversity (Tasic et al., 2016; Zeisel et al., 2015), we argue that inspection of cell type-specific marker expression should be an essential patch-seq quality control step prior to downstream analyses.

## Acknowledgements

We thank Cathryn Cadwell, Janos Fuzik, Csaba Földy, and Cedric Bardy for sharing data and Jim Berg, Dmitry Kobak, and Philipp Berens for helpful discussions. This work was supported by Kids Brain Health Network, a Canadian Institute for Health Research Post-doctoral Fellowship, and NIH grant MH111099.

## Author contributions

SJT and PP conceived the project. SJT implemented the methodology and generated the results with assistance from LT, OBM, CB and MB. All authors contributed to interpreting the results. SJT and PP wrote the paper with assistance from all authors.

## Competing interests

The authors declare no competing financial interests.

---

**Supplement Figure 1.**
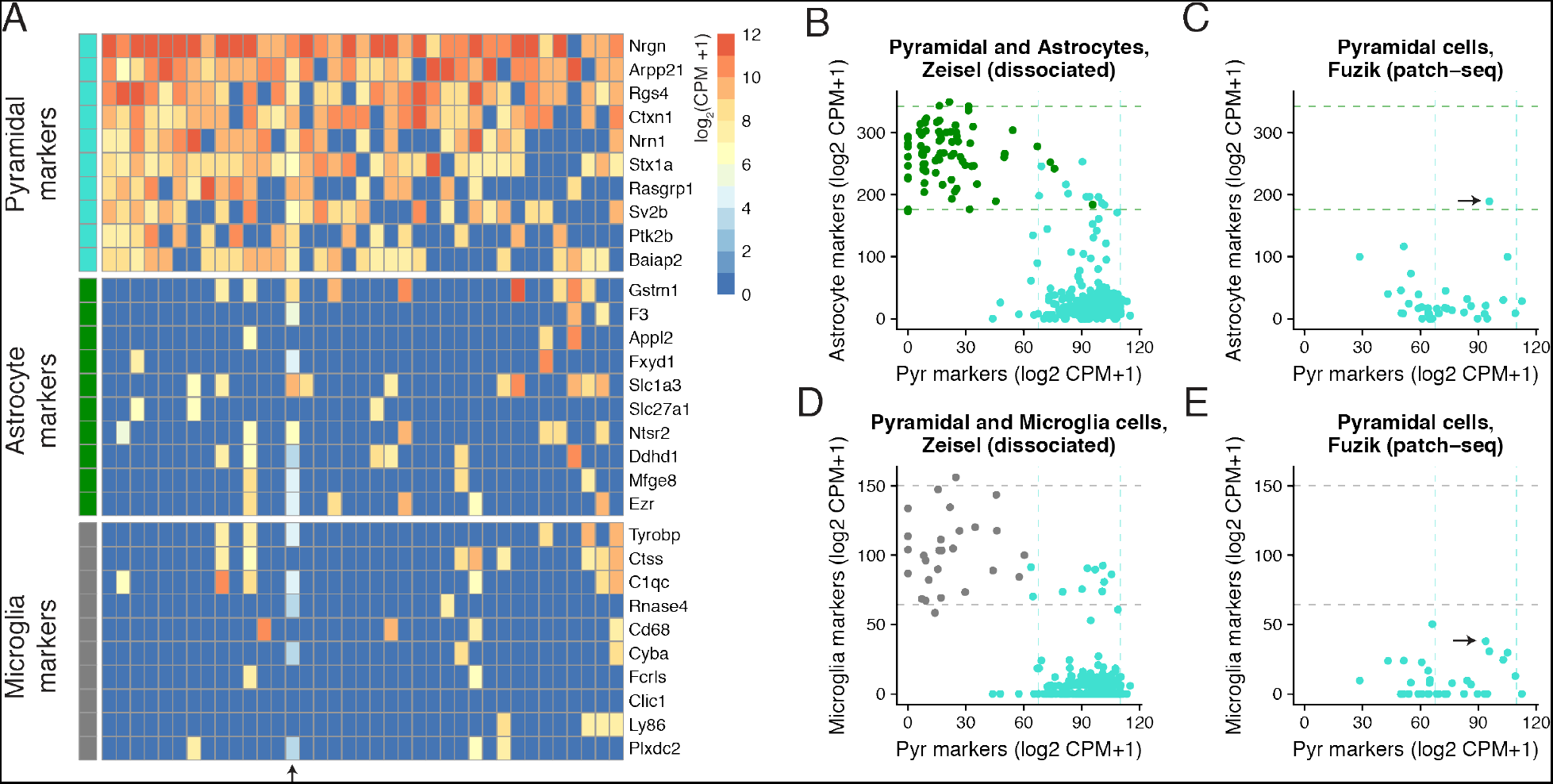
Expression of cell type-specific marker genes in patch-seq samples from Fuzik. A) Gene expression profiles for sampled pyramidal cells for various cell type-specific markers. B) Summed expression of cell type-specific marker genes for Pyramidal cell (x-axis) and Astrocyte (y-axis) markers. Dots reflect cortical Pyramidal cell (turquoise) and Astrocyte (green) single cells collected in the Zeisel dataset, based on dissociated scRNAseq. Dashed lines reflect 95% intervals of marker expression for each cell type. C) Same as B, but showing summed marker expression for Pyramidal cells shown in A based on patch-seq data. Arrow denotes the same single-cell highlighted in A. D,E) Same as B and C, but showing comparison of microglial marker expression.

**Supplement Figure 2.**
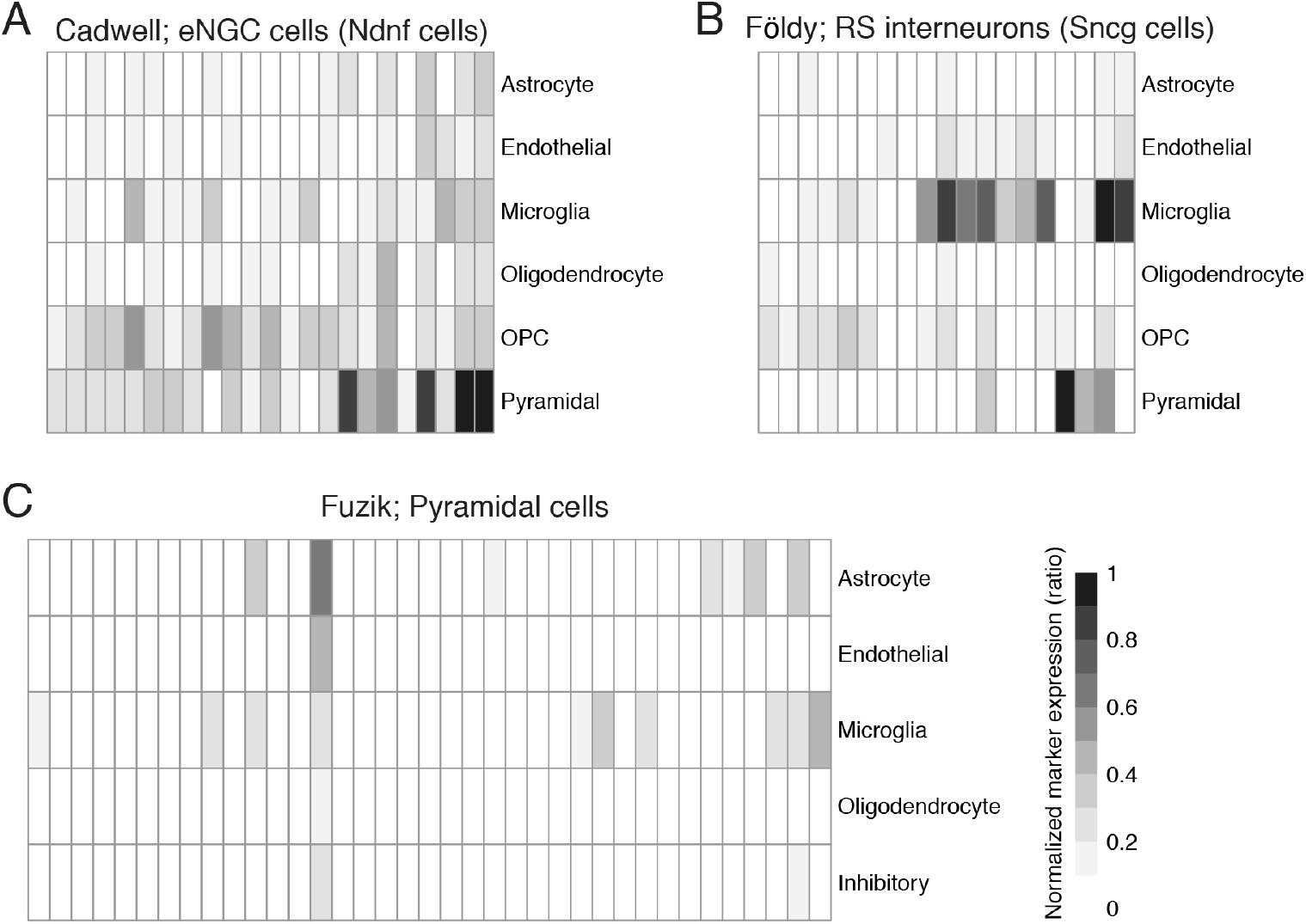
Summarized marker gene expression in patch-seq samples for broad cell classes. A) Cortical Layer 1 elongated neurogliaform cells (eNGCs) from Cadwell; B) Hippocampus regular spiking (RS) GABAergic interneurons from Földy; C) Cortical Pyramidal cells from Fuzik. Each column reflects a single-cell sample and columns are sorted as in Figure 1 and Supplement Figure 1. Heatmap colors show cell type-specific marker expression, normalized to expected expression based on dissociated cell reference datasets (Tasic, A, B; Zeisel, C). 0 indicates little-to-no detected off cell-type marker contamination (relative to dissociated cells) and 1 indicates strong expression of off-cell-type markers. Oligodendrocyte precursor cells not available in C because this cell type was not explicitly annotated in the Zeisel dataset.

**Supplement Figure 3.**
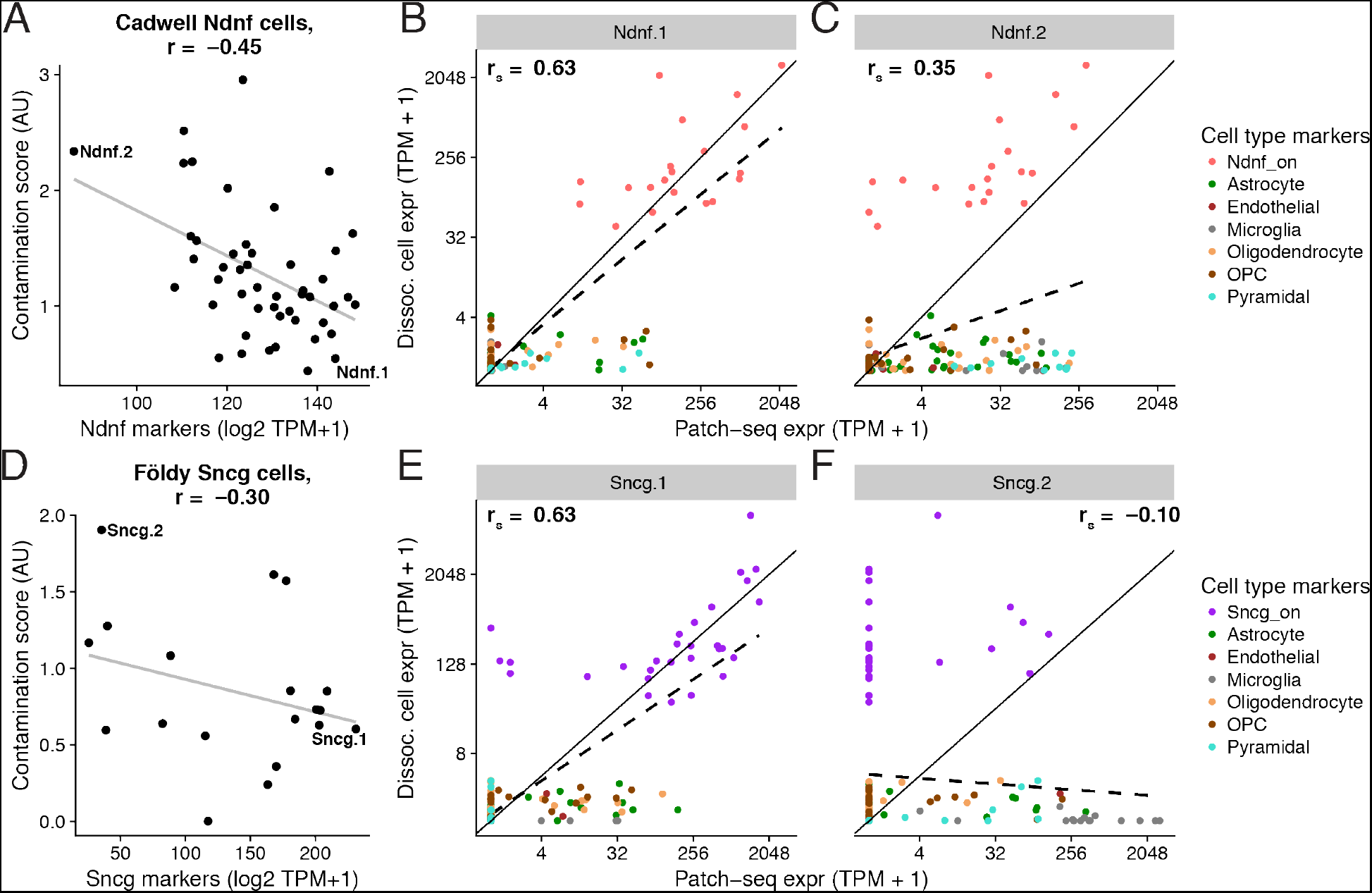
Relationship between inferred contamination and endogenous marker expression. A) Summed expression of endogenous “on”-cell type cellular markers (x-axis) versus normalized contamination indices (y-axis, summing across normalized contamination values across broad cell types) for individual Ndnf cells from the Cadwell dataset (dots). B, C) Examples of “on”- and “off”-cell type marker expression for two single-cell patch-seq samples indicated in A. X-axis shows expression of marker genes (dots) in an individual patch-seq sampled cell and y-axis shows the average expression of the same markers in Ndnf-type dissociated cells from Tasic. Solid line is unity line, dashed line shows best linear fit, and rs denotes Spearman correlation between patch-seq and mean dissociated cell marker expression. Cell Ndnf.1 (shown in B) illustrates a patch-seq sample with high expression of “on”-type endogenous markers and relatively little “off”-cell type marker expression whereas cell Ndnf.2 (shown in C) expresses endogenous markers less strongly (relative to dissociated cells of same type) and higher levels “off”-cell type marker expression. D-F) Same as A-C, but for hippocampal GABAergic regular spiking interneurons (i.e., Sncg cells) characterized in Földy dataset.

**Supplementary Table 1:** Description of dissociated-cell scRNAseq datasets and patch-clamp electrophysiological datasets used. For RNA amplification, the Tasic scRNAseq dataset employed SMARTer (i.e., Smart-seq based, consistent with the Cadwell, Foldy, and Bardy datasets) whereas the Zeisel dataset employed C1-STRT (consistent with the Fuzik dataset)

| Dataset | Experiment type | Preparation | Description | Accession | Number of cells |
| --- | --- | --- | --- | --- | --- |
| Tasic<br>(Tasic et al., 2016) | dissociated cell scRNAseq | Dissociated cells | Visual cortex neurons and glia | GSE71585 | 1366 |
| Zeisel<br>(Zeisel et al., 2015) | dissociated cell scRNAseq | Dissociated cells | Somatosensory cortex and hippocampus neurons and glia | GSE60361 | 3005 |
| Allen Institute Cell Types<br>(Teeter et al., 2018) | patch-clamp electrophysiology | Acute mouse slices | Visual cortex neurons | celltypes.brain-map.org | 952 |

**Supplementary Table 2:** Matching of patch-seq cell types to dissociated cell reference atlases.

| Patch-seq dataset | Cell type (patch-seq) | Matched cell type (dissociated cell; Tasic) | Matched cell type (dissociated cell; Zeisel) |
| --- | --- | --- | --- |
| Cadwell | Cortex Layer 1 elongated neurogliaform cell (eNGC) | Ndnf cluster (Ndnf Car4, Ndnf Cxcl14) | Int12, Int15 |
| Cadwell | Cortex Layer 1 single bouquet cells (SBC) | Ndnf cluster (Ndnf Car4, Ndnf Cxcl14) | Int12, Int15 |
| Földy | Hippocampus regular-spiking (RS) interneurons | Sncg | Int 5 |
| Földy | Hippocampus CA1 and Subiculum Pyramidal cells | Pyramidal cluster | Pyramidal cluster (excluding CA1PyrInt) |
| Földy | Hippocampus fast-spiking (FS) interneurons | Pvalb cluster (Pvalb Gpx3, Pvalb Wt1, Pvalb Tacr3, | Int 3 |
|  |  | Pvalb Tpbg, Pvalb Cpne5, Pvalb Rspo2, Pvalb Obox3) |  |
| Fuzik | Cortex Layer 1 and 2 interneurons | Ndnf cluster (Ndnf Car4, Ndnf Cxcl14) | Int12, Int15 |
| Fuzik | Cortex Pyramidal cells | Pyramidal cluster | Pyramidal cluster (excluding CA1PyrInt) |

**Supplementary Table 3.** Mapping of broad cell types between Tasic and Zeisel dissociated cell reference datasets. * denotes oligodendrocyte precursor cell type not being explicitly labelled in Zeisel.

| Broad cell type | Tasic subtypes | Zeisel subtypes |
| --- | --- | --- |
| Astrocyte | Astro Gja1 | Astro2, Astro1 |
| Endothelial | Endo Myl9, Endo Tbc1d4 | Vsmc |
| Inhibitory | Vip Chat, Vip Parm1, Vip Mybpc1, Vip Gpc3, Pvalb Gpx3, Ndnf Cxcl14, Vip Sncg, Ndnf Car4, Sst Myh8, Sst Th, Sst Chodl, Sst Tacstd2, Sst Cdk6, Pvalb Wt1, Sncg, Sst Cbln4, Pvalb Tacr3, Igtp, Smad3, Pvalb Tpbg, Pvalb Cpne5, Pvalb Rspo2, Pvalb Obox3 | Int10, Int6, Int9, Int2, Int4, Int1, Int3, Int13, Int16, Int14, Int11, Int5, Int7, Int8, Int12, Int15 |
| Microglia | Micro Ctss | Mgl1, Mgl2 |
| Oligodendrocyte | Oligo Opalin, Oligo 96_Rik | Oligo1, Oligo3, Oligo4, Oligo2, Oligo6, Oligo5 |
| OPC | OPC Pdgfra | * |
| Pyramidal | L2/3 Ptgs2, L2 Ngb, L4 Ctxn3, L4 Scnn1a, L5a Batf3, L5a Pde1c, L6a Mgp, L6b Serpinb11, L6b Rgs12, L5a Hsd11b1, L4 Arf5, L5a Tcerg1l, L6a Sla, L6a Syt17, L6a Car12, L5b Cdh13, L5 Ucma, L5b Tph2, L5 Chrna6 | S1PyrL4, ClauPyr, S1PyrL5, S1PyrL23, S1PyrDL, S1PyrL5a, SubPyr, CA1Pyr1, S1PyrL6b, S1PyrL6, CA1Pyr2, CA2Pyr2 |

Supplementary Table 4: List of cell type-specific markers based on re-analysis of published dissociated cell-based scRNAseq experiments from mouse brain.

